# Variation in whole-body regeneration between *Botrylloides* morphs and species

**DOI:** 10.1101/2022.07.12.499812

**Authors:** Berivan Temiz, Megan J. Wilson

## Abstract

Regeneration is a characteristic of the animal kingdom, and regenerative capacity is limited to higher taxonomical levels. In contrast, some basal animals like urochordates maintain a unique regeneration capacity, such as undergoing whole-body regeneration (WBR), an ability not observed in other chordates. Botryllids are colonial urochordates that can recreate new bodies through WBR from solely vascular tissue within ^~^2 weeks. To date, some species from the botryllid family were reported to orchestrate WBR, including *B. diegensis*. This study provided two novel records of WBR of *B. jacksonianum* and *B. aff. anceps* along with the two distinct morphs of *B. diegensis*. Interestingly, *B. aff. anceps* executed twin-body regeneration while this was limited to one-body for *B. jacksonianum* and *B. diegensis*. Histological sections validate the formation of multiple niches during WBR. Furthermore, the process of regeneration is phenotypically more similar between *B. aff. anceps* and *B. diegensis*. In contrast, the type of WBR in *B. jacksonianum* is similar to vascular budding as the niches were built from the vascular epithelium without undergoing significant tissue remodelling.

## 1. INTRODUCTION

All animals have the capacity to regenerate tissues/cells to some extent. However, substantial regenerative ability (whole organs and structures) predominantly diminishes during evolutionary development (Fig. 1A) (Poss, 2010). The capacity to regenerate varies among the animal phylum and in different tissues, which often negatively correlates with the tissue and body complexity (Zhao et al., 2016). Furthermore, this ability usually diminishes during maturation, particularly after metamorphosis or puberty (Murawala & Knapp, 2021). While the enhanced investment in this ability, such as regeneration of the whole adult body, is an advantage for some animals’ survival, it can result in poor investment in other resources such as sexual reproduction that may increase the organism’s fitness (Bely & Nyberg, 2010; Grillo et al., 2016).

**Figure 1.**
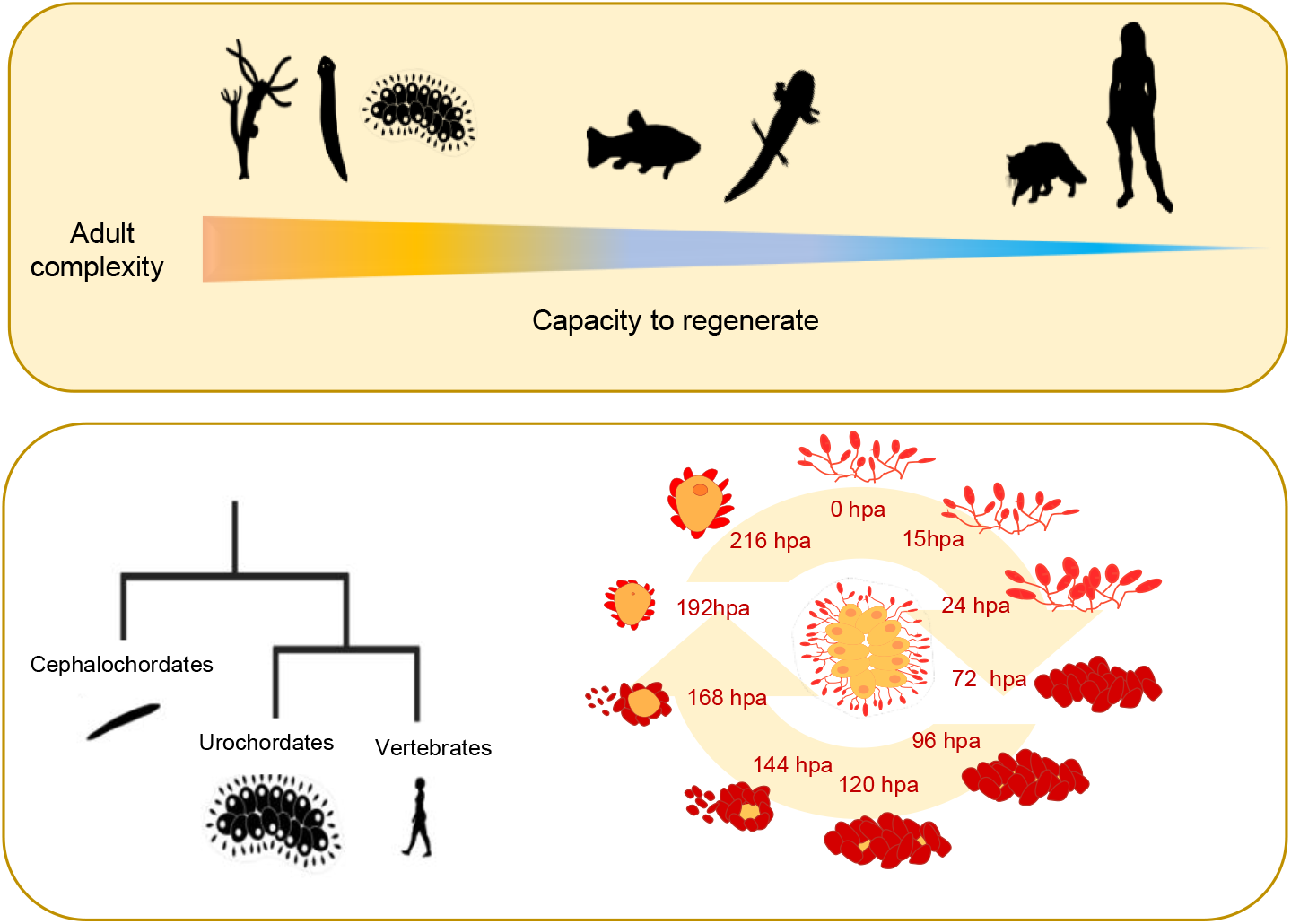
**A) Regeneration capacity generally diminishes with complexity within the animal kingdom.** The colour distribution indicates the spectrum of regenerative capacity while highest in invertebrates such as hydra, planaria and tunicate species and decreasing towards the mammals. **B) Phylogenetic proximity of urochordates to vertebrates.** The tree represents the phylogenetic position of tunicates as the closest chordate group to the vertebrates. **C) Whole body regeneration of *B. diegensis*.** The process of *B. diegensis* whole body regeneration in the clockwise direction. WBR is initiated from a vascular fragment depleted from zooids following the contraction of ampullae. Regeneration niches are located within the condensed tissue, where the zooid starts to form. WBR can be completed in as little as 216 hours.

Whole body regeneration (WBR) is the restoration of a complete functional individual from a remaining piece of tissue. This phenomenon is rare in chordate animals and restricted to some invertebrate groups. The phylum Chordata can be divided into three sub-phyla as Tunicata (Urochordata), Cephalochordata and Vertebrata (Fig. 1B). Within the chordates, one of the most amazing regenerative abilities is seen in teleost fish and urodele amphibians which can renew their tissues, body fragments and organs. However, mammals have a limited regeneration capacity. While organs like the liver can be renewed, most tissues, such as digit tips, can only be restored during a particular life stage, as in mice (Han et al., 2003). In contrast, WBR is limited to some groups like urochordates and has never been recorded in vertebrates (Blanchoud et al., 2017). Therefore, as the closest relative to vertebrates and highly regenerative animals, urochordates are a popular group for regeneration studies and understanding chordate evolution.

*B. diegensis* is a sessile colonial tunicate that demonstrates the clearest example of WBR within the botryllids. This species can propagate WBR from a small fragment of vascular tissue in 8 to 14 days, and as few as 100 cells are enough for rebuilding a functional body (Fig. 1C) (Rinkevich et al., 1995; Rinkevich et al., 2007a; Zondag et al., 2016; Blanchoud et al., 2017). WBR is triggered after the loss of all individuals in the system after a short healing response to cease haemolymph diminution and is initiated through stem-like cells located in the vascular tissue (Rinkevich et al., 2010; Blanchoud et al., 2017). The vascular termini, also called ampullae that remain in the amputated tissue, start to compact after losing all the zooids and reorganize as a composition of tiny blood cell clusters. As a result, this tissue is denser and condensed than the zooid, including colonial tissues. During WBR, blood flow is sustained through the ampullar contractions, through the heart’s pulsation in a typical colony (Blanchoud et al., 2017). Following compaction, the regeneration niches emerge within the vasculature, and one of the niches transforms into a complete zooid while others are absorbed.

Although included in different taxonomic clusters, stem cell fate during WBR of three invertebrate species, *Hydra, Planaria* and *Botrylloides*, indicate a similar characterization of reprogramming of regeneration potency that enables the conversion of cell type while maintaining the pluripotency (Knapp & Tanaka, 2012). In hydra, overexpression of a POU protein named Polynem in epithelial cells gives rise to stem cells that later express conserved stem and germ cell markers such as *Vasa, Nanos, Piwi* and *Myc* (Millane et al., 2011). Planaria sustains regeneration through the pluripotent neoblast cells and, in one particular planaria species, *Schmidtea mediterranea*, these cells express *Smed-soxP-3*, which is essential for the expansion during renewal and also homeostasis (Wagner et al., 2012). Several ancient regeneration pathways among metazoans, such as RA, NOTCH and WNT, are highly conserved in *Botrylloides* (Blanchoud et al., 2018b). *Botrylloides* WBR is predicted to be initiated through the *Piwi* expressing cells that are previously dormant in the vascular epithelium (Rinkevich et al., 2010) or through Integrin Subunit Alpha 6 (ITGA6)-positive cells in the circulation (Kassmer et al., 2020).

*B. diegensis* is unique in the way it undergoes WBR. Unlike other animals undergoing epimorphic regeneration, it does not form a blastema. Instead, it forms a regeneration niche inside the vasculature. *B. diegensis* initiates regeneration at multiple sites, while in other species, a single local induction signal results in regeneration at a single site (Rinkevich et al., 2007b). Although many regeneration niches are occurring, only the fastest developing one completes the regeneration of *B. diegensis* by developing into an adult zooid that undergoes organogenesis. The other buds undergo a degeneration process by apoptosis and are absorbed into the adult zooid and the surrounding vasculature (Rinkevich et al., 2007b).

Although several botryllids are known to undergo vascular budding as a part of the asexual cycle that is phenotypically similar to WBR, this happens when zooids are still present in the colonial system, unlike WBR (Oka & Watanabe, 1959; Saito & Watanabe, 1985; Okuyama & Saito, 2001). Nevertheless, the WBR of *B. diegensis* and *Botrylloides violaceus* is orchestrated solely without any zooid, bud or budlet tissue present (Rinkevich et al., 1995; Brown et al., 2009). In contrast to the WBR of *B. diegensis*, multiple zooids are formed in the WBR of *B. violaceus*. Recently, *B. anceps* was reported to orchestrate single zooid WBR via stem-cell niche formation, while there is no previous record on *B. jacksonianum* (Karahan et al., 2022).

Despite the morphological and anatomical insights, the cellular origin of botryllid WBR is still enigmatic. Some mechanisms of the WBR are equivalent to those in the asexual cycle and embryogenesis due to similarity in the expression of the markers such as *Piwi, Vasa* and *Sox* (Lai & Aboobaker, 2018). However, characterization of the progenitor cells that initiate WBR requires a more profound molecular identification. In addition, variations in regenerative capacity between sister species might explain the execution of WBR at the molecular level.

## 2. MATERIALS & METHODS

### 2.1. Animal husbandry & Regeneration Assay

Colonies were sampled from the Otago Harbour (45°52’17’’S, 170°31’43’’E) of New Zealand between June 2020 and April 2022 from submerged parts of ropes, pontoons, and tires. The colonies were attached to 5 x 7.5-cm glass slides and placed horizontally for 1 day in 200 ml of still seawater. The colonies were taken to the tanks the next day and set vertically in still seawater. The following day, tanks started to be aerated, and the colonies were fed. Their water was changed with filtered seawater every two days and provided a shellfish diet (LPB Frozen Shellfish Diet™ – Reed Mariculture). Finally, the slides were cleaned with a paintbrush to avoid algal overgrowth.

Different morphs and species of *Botrylloides* were checked to determine whether they could undergo WBR. All zooids, buds, and budlets were ablated from the colonies using a scalpel and a single-edged razor blade, leaving only the vascular tissue. The slides were placed back in aerated saltwater tanks and monitored for daily regeneration.

### 2.2. Histology

Regenerating colonies were fixed in 4% paraformaldehyde for 2 h and embedded in paraffin wax. The tissues were sectioned for 5 μm thickness in the transverse or longitudinal plane. Hematoxylin and eosin (H&E) stains were applied to the tissue sections deparaffinized with xylene and rehydrated with serial dilutions of ethanol. H&E staining was followed as 1) Staining for 4 min in hematoxylin 2) Washing for 2 min in tap water 3) 2 min in Scott’s tap water (2 g potassium bicarbonate and 20 g MgSO_4_ /L) 4) Tap water 2 min 5) Eosin staining for 30 s 6). Slides were dehydrated with ethanol, then xylene, and coverslipped. Images were taken using a Nikon Ti2E Inverted Brightfield microscope.

## 3. RESULTS

Regeneration of different botryllid morphs and species was monitored after vascular fragments from healthy colonies were amputated (no zooids, buds or budlets present, and the colony is not during the take-over phase). The brown-orange morph of *B. diegensis* regenerated a functional zooid after 12 days post-amputation (dpa) (Fig. 2A). The blood tissue compacted during the early stages of the WBR. A visible regeneration niche was observed from 6 dpa and started to form, where the development of the zooid was seen in the later stages. Likewise, the purple-white (or brown-white) morph of *B. diegensis* regenerated a healthy zooid after 13 dpa (Fig. 2B). Due to the low contrast by the pigment cells, the niche opening could be observed during 9 dpa.

**Figure 2.**
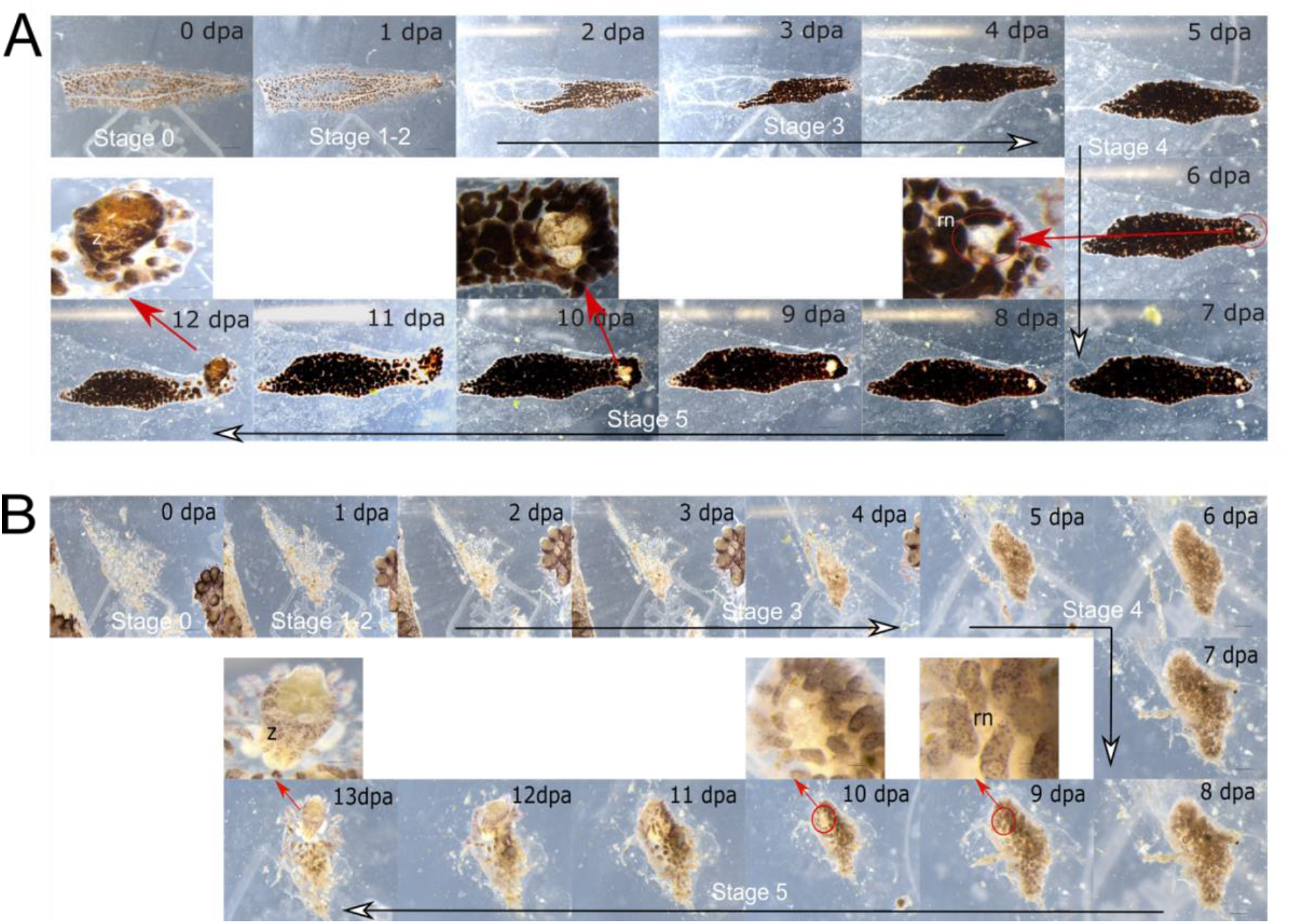
WBR of two morphs of *Botrylloides diegensis*. **A**) WBR of *B. diegensis* brown-orange morph. After 6 days, a regeneration niche is visible. By 12 days, a filter-feeding adult is present. **B**) WBR of *B. diegensis* purple-white morph. Following compaction of the vascular tissue, a regeneration niche is observed at 9 dpa. Organogenesis is completed by 12 dpa. dpa: day post-amputation, rn: regeneration niche, z: zooid. Stage 0 (0 hpa), stage 1 (15 hpa), stage 2 (24 hpa), stage 3 (2 - 4 dpa), stage 4 (5 - 7 dpa), stage 5 (8 to 13 dpa). White arrowheads show the continuity of the regeneration stage.

The two morphs of *B. diegensis* share identical stages of regeneration, which were previously identified for the orange morph of *B. diegensis* (Blanchoud et al., 2017) (Fig. 2). Stage 0 (0 hpa) vascular tunic at injury, the blood flow slows within minutes to reduce haemolymph loss (Fig. 2A-B). The cut sites start to heal during stage 1 (15 hpa). Stage 2 (24 hpa) involves remodelling the terminal ampullae, which change shape towards a more spheroidal formation. The vascular tissue becomes a compact blood mass during stage 3 (2 - 4 dpa). By stage 4 (5 - 7 dpa), regeneration niches emerge and can be observed as clear-like vesicles. In the final stage (Stage 5 - 8 to 13 dpa), organogenesis starts, and regeneration is completed resulting in a single functional zooid (Fig. 2A-B).

Like *B. diegensis*, an amputated vascular fragment of *B. jacksonianum* developed a single zooid after 15 dpa (Fig. 3A). The tissue did not undergo compaction or vascular remodelling as observed with *B. diegensis* and *B. aff. anceps* (Fig. 2 and 3B). Two regeneration niches emerged; only one completed regeneration of a zooid, visible after 8 dpa (Fig. 3A). The colony reabsorbed the other niche. Regeneration was completed in 15 days.

**Figure 3.**
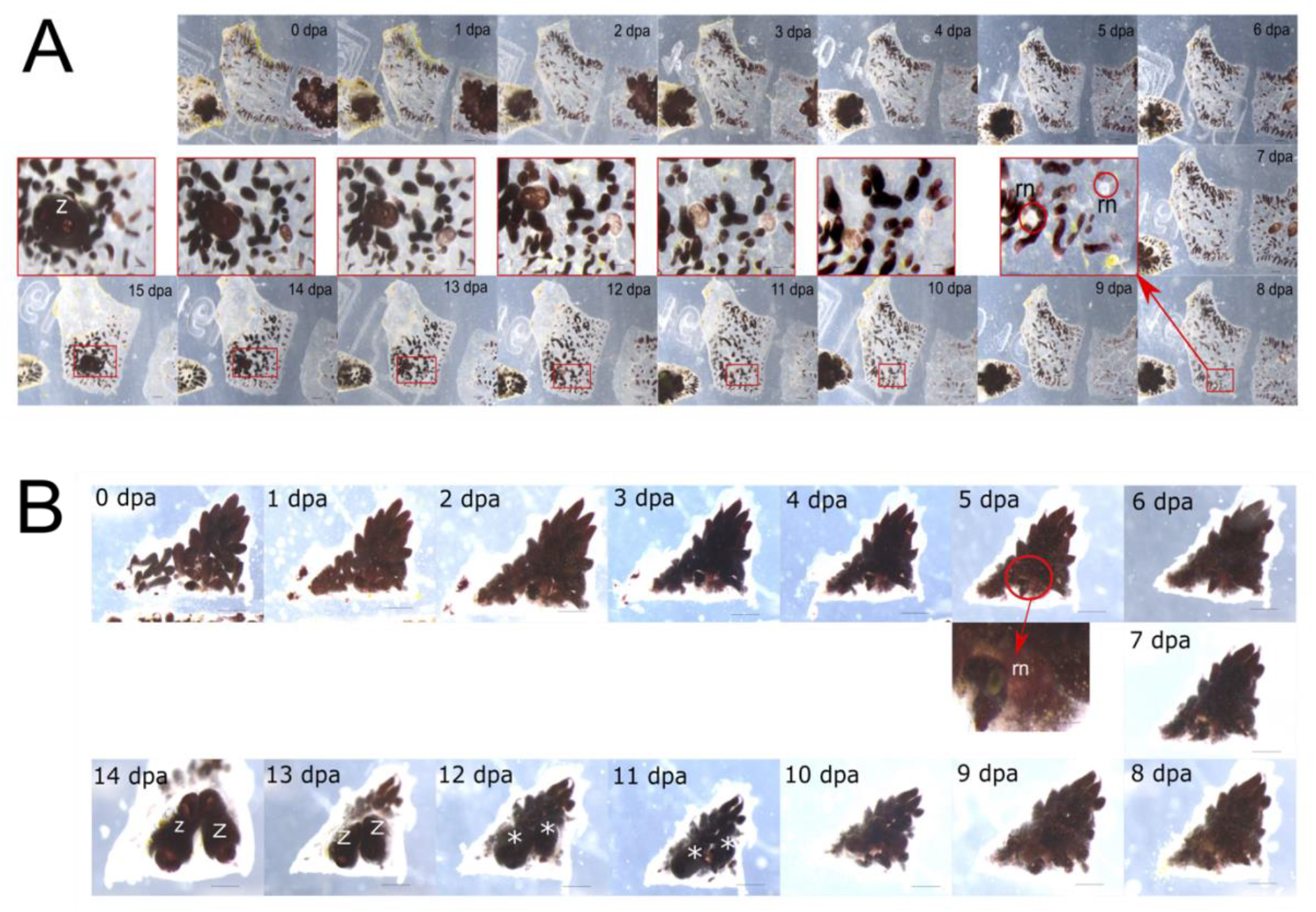
WBR of *Botrylloides jacksonianum* and *Botrylloides affinis anceps*. **A**) WBR of *B. jacksonianum*. The regeneration takes place in 15 dpa. Niches were first observed at 8 dpa. **B**) *B. aff. anceps* can create two functional zooids after 15 dpa. Niche is seen after 5 dpa. dpa: day post-amputation, rn: regeneration niche, z: zooid.

In contrast, *B. aff. anceps* created two fully functional individuals after 14 dpa (Fig. 3B). The niche was visible since 5 dpa. While the type of niche formation and duration resembled the *B. diegensis*, more than one niche could complete the organogenesis resulting in two zooids forming simultaneously. We validated the regeneration of the *Botrylloides* species and morphs by cutting multiple fragments (Table 1). While *B. diegensis* and *B. jacksonianum* always created a single zooid, *B. aff. anceps* produced double zooids more than 50 % of the time.

**Table 1.**
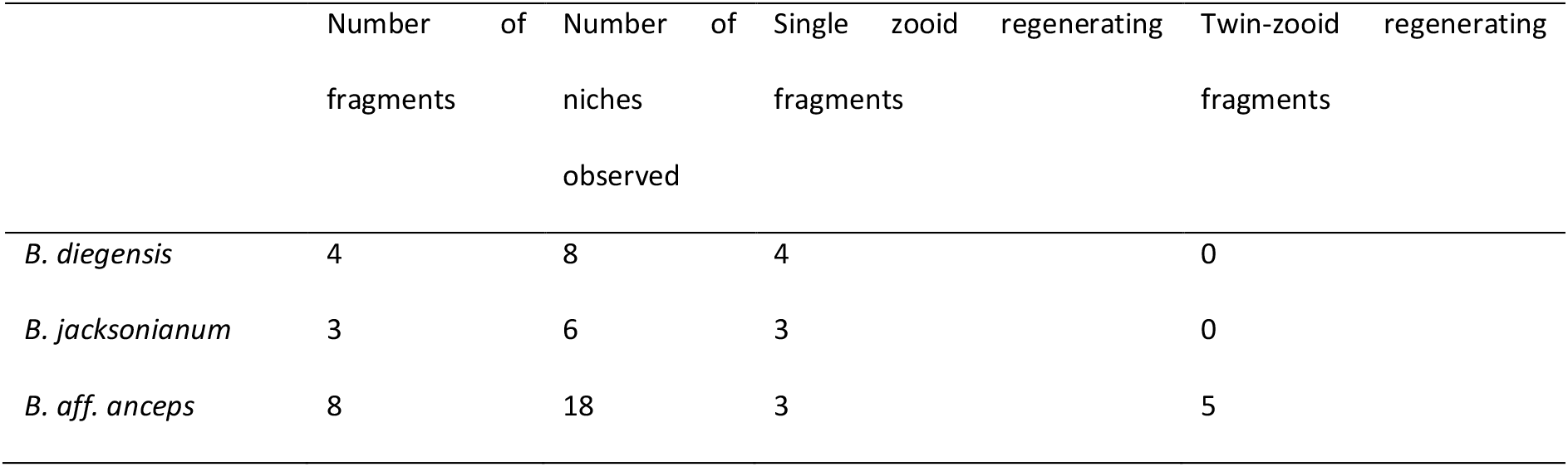
Regeneration assay of *B. diegensis, B. jacksonianum* and *B. aff. anceps*

Histological examination of *B. aff. anceps* and *B. jacksonianum* WBR revealed a niche formation was similar to that of *B. diegensis* (Blanchoud et al., 2017) (Fig. 4). For *B. aff. anceps*, three regeneration niches were observed after 5 dpa (Fig. 4A and 4B). Regenerating tissue of *B. jacksonianum* had two niches beside each other during 8 dpa (Fig. 4C and 4D). Both species have similar epithelial buds forming within the niches (asterisks, Fig. 4A-D). Stem-like cells were located near the forming buds (blue arrowhead; Fig. 4B and 4D). Additionally, we observed the ubiquitous presence of macrophage-like cells in the regenerating tissue sections (white arrowheads; Fig. 4). The larger buds for *B. jacksonianum*, which have started to shape into a curved spheroid, also highlights that the tissue was at the middle or late stage 4, while *B. aff. anceps* niches were at the beginning of stage 4. We also observed fragments of vascular epithelium next to the niches (black arrowheads, Fig. 4D). These maybe derived from the thin endothelium that lines the blood vessels, although the cells appear larger.

**Figure 4.**
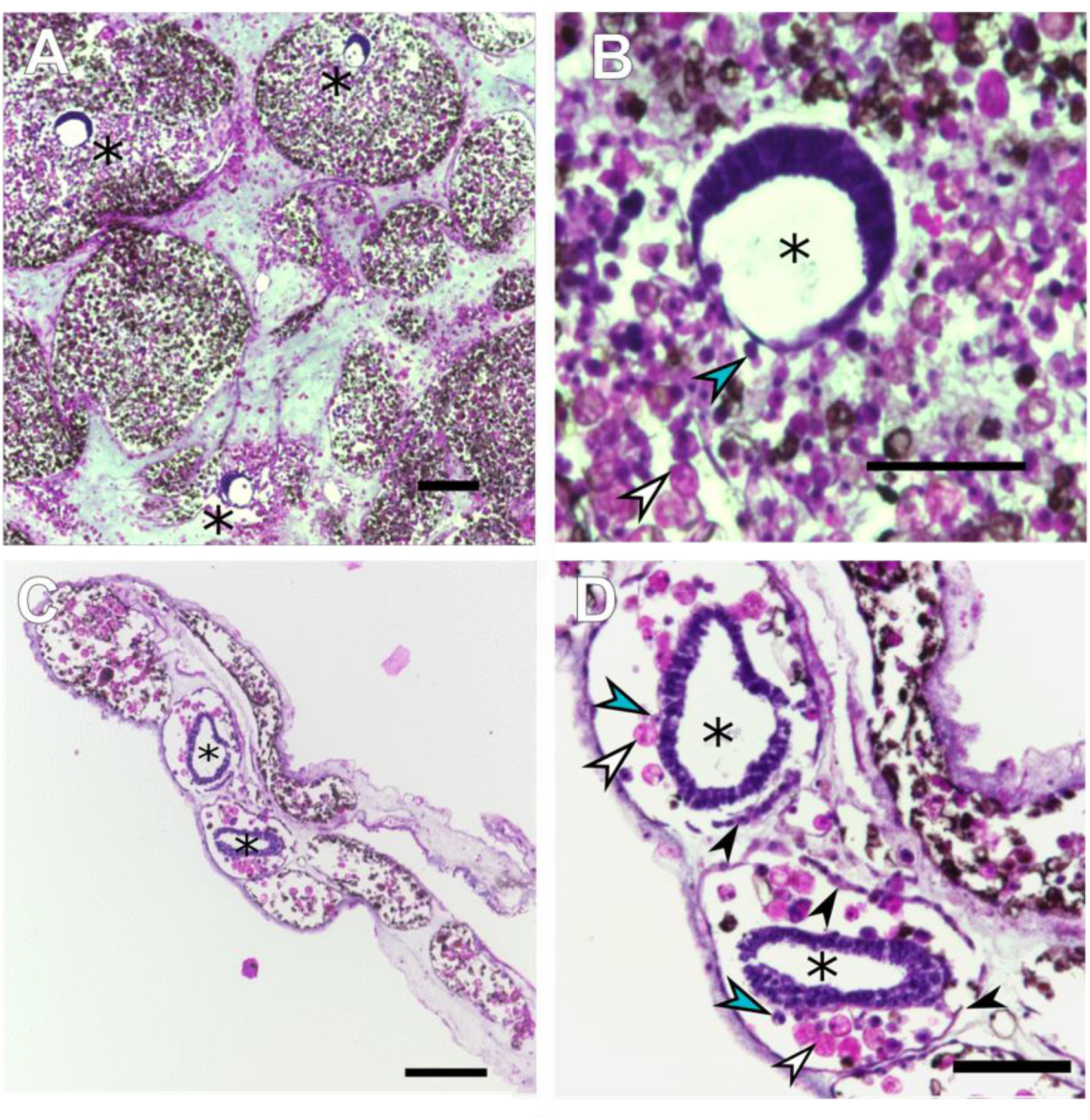
Histological examination of *B. aff. anceps* and *B. jacksonianum* during the middle stages of WBR. Sections are stained with haematoxylin and eosin. **A**) Transverse section of *B. aff. anceps* during 138 hpa (5 - 6 dpa) in lower magnification with three regeneration niches (arrowheads) **B**) Regeneration niche with *B. aff. anceps* bud in higher magnification **D**) Longitudinal section of *B. jacksonianum* during 8 dpa **D**) Two niches of *B. jacksonianum* side to side in higher magnification. Black arrowheads highlight fragmented epithelium. Blue arrowheads indicate stem cells (hemoblasts). Asterisks indicate the regeneration buds. White arrowheads point to macrophage-like cells. Scale bars are equal to 100 μm.

## 4. DISCUSSION

Botryllids share a similar life cycle, morphology, and ability to create new zooids through different developmental or regenerative programs. Recreation of the zooids in a weekly process (blastogenesis/palleal budding) is present as a part of colonial life form. In palleal budding, the primary bud develops from the position of the mature zooid, while in vascular budding, a new zooid is produced from a piece of the vascular epithelium (Oka & Watanabe, 1957). *Botrylloides lenis, Botryllus delicates* and *Botryllus primigenus* can rebuild new zooids via vascular budding (Oka & Watanabe, 1959; Saito & Watanabe, 1985; Okuyama & Saito, 2001).

Budding resembles regeneration phenotypically and involves parallel pathways (Rinkevich et al., 2010; Manni et al., 2019; Kowarsky et al., 2021). Buds are morphologically similar to the regeneration buds, including the type of cell that structures: small, round cells with dense nuclei (Oka & Watanabe, 1959; Blanchoud et al., 2017). In comparison, WBR differs from these two developmental budding processes and is only seen in the case where no zooid, bud, or budlet tissues are present. Pieces of vascular tissues of *B. jacksonianum, B. aff. anceps* and different morphs of *B. diegensis* were cut and monitored to see whether they regenerate and how different from each other. The loss of this capability can be used as a significant benchmark in searching for the cellular origin of WBR. All monitored colonies can undergo WBR in a relatively similar time course (~2 weeks), compatible with the formerly reported orange *B. diegensis* colonies (Zondag et al., 2016; Blanchoud et al., 2017). The phenotypical similarity of WBR is closer between *B. diegensis* and *B. aff. anceps* as the niches are covered in between the blood masses. In contrast, the niches of *B. jacksonianum* were visible since there was no denser blood tissue compaction. Its blood vessels were not remodelled, and the blood flow did not thicken but continued in its normal bidirectional flow.

In addition, the regeneration niches of *B. jacksonianum* were observed slightly later (7-8 dpa) than *B. aff. anceps* and *B. diegensis* (5-6 dpa), developed from the vascular epithelium, similar to vascular budding. Both B. *jacksonianum* niches are located in between a layer of epithelium. The cells that shape these two niches are also slightly different from those of *B. aff. anceps*. While almost no cytosolic compartment is present in the niche cells of *B. aff. anceps*, the ones in the *B. jacksonianum* include more cytosol and a dense nucleus similar to vascular bud cells of Oka & Watanabe (Oka & Watanabe, 1959). The regeneration niches of *B. aff. anceps* are like those of *B. diegensis*, emerging inside the ampullae. The compaction of blood tissue leaves almost no blood vessel in the system visible in the sections. Similar to the middle stages of *B. diegensis* WBR, more phagocytic cells are present around the blood tissue of both species, which have been stated to have the strong complementation of immunological pathways with the cellular differentiation events (Blanchoud et al., 2017). These cells provide removal of debris and are thought to supply nourishment during zooid development while reabsorbing the incompetent regeneration niches (Lauzon et al., 2013; Blanchoud et al., 2017).

When considering the type of niche formation, number and duration, the WBR of *B. aff. anceps* is similar to the one of *B. diegensis* but separated significantly by the twin-body regeneration. Although multiple regeneration niches can be present during the early stages of regeneration, only one single zooid completes WBR for *B. diegensis* (Rinkevich et al., 1995; Blanchoud et al., 2017; Zondag et al., 2019). This might result from a difference in regeneration potency between the sister species, nevertheless emphasizing the conservation of the regeneration mechanisms although variation is present (Blanchoud et al., 2018a; Blanchoud et al., 2018b).

## ACKNOWLEDGEMENTS

We would like to thank Robert Porteous and Vicky Clark from the University of Otago Histology team. In addition, we thank Dr. Michael Meier for the help during samplings. This study was supported by the Royal Society of New Zealand Marsden fund grant (UOO1713). B Temiz was supported by a Department of Anatomy, University of Otago PhD scholarship.

## DATA AVAILABILITY STATEMENT

The data supporting these findings are openly available.

